# Pore-forming activity of *S. pneumoniae* pneumolysin disrupts the paracellular localization of the epithelial adherens junction protein E-cadherin

**DOI:** 10.1101/2023.06.01.543286

**Authors:** Shuying Xu, Devons Mo, Fatima Z. Rizvi, Juan P. Rosa, Jorge Ruiz, Shumin Tan, Rodney K. Tweten, John M. Leong, Walter Adams

**Affiliations:** Department of Molecular Biology and Microbiology, Tufts University, Boston, MA, 02111 USA; Program in Immunology, Tufts Graduate School of Biomedical Sciences, Boston, MA, USA; Department of Biological Sciences, San Jose State University, San Jose, CA 95192 USA; Department of Microbiology and Environmental Toxicology, University of California, Santa Cruz, Santa Cruz, CA, USA; University of Puerto Rico, Cayey, PR 00737 USA; Francisco de Vitoria University, Madrid, Spain; Department of Microbiology and Immunology, University of Oklahoma Health Sciences Center, Oklahoma City, OK 73104 USA; Stuart B. Levy Center for Integrated Management of Antimicrobial Resistance at Tufts (Levy CIMAR), Boston, Massachusetts, USA

**Keywords:** neutrophils, epithelial cell junction, E-cadherin, airway mucosal barrier, cholesterol-dependent cytolysin

## Abstract

*Streptococcus pneumoniae*, a common cause of community-acquired bacterial pneumonia, can cross the respiratory epithelial barrier to cause lethal septicemia and meningitis. *S. pneumoniae* pore-forming toxin pneumolysin (PLY) triggers robust neutrophil (PMN) infiltration that promotes bacterial transepithelial migration *in vitro* and disseminated disease in mice. Apical infection of polarized respiratory epithelial monolayers by *S. pneumoniae* at a multiplicity of infection (MOI) of 20 resulted in recruitment of PMNs, loss of 50% of the monolayer, and PMN-dependent bacterial translocation. Reducing the MOI to 2 decreased PMN recruitment two-fold and preserved the monolayer, but apical-to-basolateral translocation of *S. pneumoniae* remained relatively efficient. At both MOI of 2 and 20, PLY was required for maximal PMN recruitment and bacterial translocation. Co-infection by wild type *S. pneumoniae* restored translocation by a PLY-deficient mutant, indicating that PLY can act in *trans*. Investigating the contribution of *S. pneumoniae* infection on apical junction complexes in the absence of PMN transmigration, we found that *S. pneumoniae* infection triggered the cleavage and mislocalization of the adherens junction (AJ) protein E-cadherin. This disruption was PLY-dependent at MOI of 2 and was recapitulated by purified PLY, requiring its pore-forming activity. In contrast, at MOI of 20, E-cadherin disruption was independent of PLY, indicating that *S. pneumoniae* encodes multiple means to disrupt epithelial integrity. This disruption was insufficient to promote bacterial translocation in the absence of PMNs. Thus, *S. pneumoniae* triggers cleavage and mislocalization of E-cadherin through PLY-dependent and PLY-independent mechanisms, but maximal bacterial translocation across epithelial monolayers requires PLY-dependent neutrophil transmigration.

## Introduction

*Streptococcus pneumoniae* (the pneumococcus, or SPN) is the most frequently isolated causative pathogen of pneumonia, with over 150,000 hospitalizations annually in the U.S. alone and a mortality rate of 5-7% (1). An ominous development in the course of pneumococcal disease is the translocation of bacteria from the lung into the bloodstream. Indeed, the case fatality rate of pneumococcal disease increases to 15-20% upon the development of bacteremia (2). The respiratory epithelium, like other mucosal surfaces, serves as a first line of defense against invading pathogens by maintaining a physical barrier comprised of tight junction (TJ) and adherens junction (AJ) multiprotein complexes. (3). TJ proteins, such as scaffolding proteins zonula occludens (ZO) and transmembrane proteins claudins, occludins, and junctional adhesion molecules (JAM), localize to the apical region of cell-cell junctions (4) where they form an intercellular membrane fence (5). AJ proteins, such as E-cadherin and catenins, are located basolateral to TJs and are essential for the formation and maturation of cell-cell contacts (6). Orchestration of junction protein abundance, localization, coupling to the cytoskeleton, and interaction with multiple cellular signaling pathways are all essential for effective epithelial barrier function (3).

Models for pneumonia and for other pulmonary diseases such as acute lung injury, asthma, and acute respiratory distress syndrome feature loss of pulmonary epithelial integrity and concomitant deleterious outcome (7, 8). Indeed, *S. pneumoniae* infection reduces epithelial junction organization in explanted human lung tissue (4), and murine models suggest that *S. pneumoniae* modulates airway epithelial junctions, influencing the degree of paracellular bacterial migration (9, 10). *S. pneumoniae* pneumolysin (PLY), a major *S. pneumoniae* virulence factor that belongs to the of cholesterol-dependent cytolysins (CDCs) that form ∼25 nm diameter pores in mammalian cell membranes, may play a critical role in this process. Pore formation by CDCs activates cytoskeletal rearrangement (11) and calcium flux (12), two processes that can lead to the breakdown of cellular junctions. Two separate studies have implicated PLY in damaging TJs (13, 14), such as ZO-1. In addition, other *S. pneumoniae* factors have been reported to downregulate tight junction components, such as TLR stimulants (9) and pneumococcal neuraminidase (15).

A hallmark of *S. pneumoniae* lung infection is the massive influx of neutrophils (polymorphonuclear leukocytes, or PMNs) into airspaces (16, 17), and in addition to directly disrupting epithelial integrity, PLY is a potent activator of inflammation. PLY triggers the production of neutrophil chemoattractants (18, 19), including epithelial production of the eicosanoid hepoxilin A3 (HXA_3_), resulting in PMN transepithelial migration (20, 21). Intranasal and intratracheal challenge of mice with *S. pneumoniae* has demonstrated an important role for PMNs in the clearance of pneumococci early in the infection (17, 22), in part reflecting their diverse range of anti-pneumococcal effector mechanisms such as the production of reactive oxygen species (ROS), proteases and antimicrobial peptides, and neutrophil extracellular traps (17, 23–25). However, these same effector molecules can damage intercellular junctions and increase epithelial permeability (26–28), with prolonged inflammatory PMN responses leading to epithelial barrier disruption, pulmonary edema, and significant lung damage (29, 30). In some mouse models, retention of high numbers of pulmonary PMNs after the initial acute infection stage is associated with bacteremia and lethality, indicating that inappropriate PMN responses exacerbate pneumococcal disease (21, 22). For example, upon pneumococcal lung challenge in mice, diminishing PMN pulmonary infiltration by blocking HXA_3_ production effectively decreased bacteremia and promoted 100% survival of an otherwise uniformly fatal infection (21, 31).

The impact of these diverse PLY functions is reflected in the observation that in several pulmonary infection mouse models, PLY is required for maximal inflammation, tissue damage, and bacteremia (32–34). Although the majority of the known effects of PLY on host cells is linked to PLY’s pore-forming activity (35), PLY interacts with non-cholesterol receptors such as the mannose receptor C type 1 (MRC-1) on macrophages and dendritic cells (36) and is capable of altering functions of both immune and non-immune cells at sub-cytolytic concentrations (37). In addition, PLY-deficient strains are capable of causing bacteremia (38), suggesting that *S. pneumoniae* encodes multiple factors that contribute to bacterial dissemination. Here, we subjected polarized respiratory epithelial monolayers to apical infection by *S. pneumoniae* and systematically examined the role of infectious dose, PLY-mediated pore formation, and PMN transmigration on the integrity and organization of the AJ protein E-cadherin, as well as on bacterial translocation.

## Materials and Methods

### Bacterial strains and growth conditions

Mid-exponential growth phase aliquots of *S. pneumoniae* TIGR4 (serotype 4) were grown in Todd-Hewitt broth (BD Biosciences) supplemented with 0.5% yeast extract in 5% CO_2_ and Oxyrase (Oxyrase, Mansfield, OH), and frozen in growth media with 20% (v/v) glycerol. Bacterial titers in aliquots were confirmed by plating serial dilutions on Tryptic Soy Agar plates supplemented with 5% sheep blood agar (Northeast Laboratory Services, Winslow, ME). The TIGR4 pneumolysin-deficient mutant (Δ*ply*) and GFP-expressing WT and Δ*ply* TIGR4 were gifts from Dr. Andrew Camilli (Tufts University, Boston, MA). The cholesterol-binding activity-deficient pneumolysin mutant (CBS) was generated by transformation of wild type TIGR4 with linear CBS DNA amplified from *Escherichia coli* plasmid gifted by Dr. Rod Tweten (University of Oklahoma, OH), and screened for non-hemolytic transformants. Insertion of the CBS *ply* gene at the original locus was confirmed by sequencing. For experiments, *S. pneumoniae* strains were grown in Todd-Hewitt broth (BD Biosciences), supplemented with 0.5% yeast extract and Oxyrase, in 5% CO_2_ at 37 °C and used at mid-log phase to late log phase.

### Growth and maintenance of epithelial cells

Human pulmonary mucoepidermoid carcinoma-derived NCI-H292 (H292) cells were grown on the underside of collagen-coated Transwell filters (5 µm pore size, 0.33-cm^2^, Corning Life Sciences) in RPMI 1640 medium (American Type Culture Collection, Manassas, VA) supplemented with 2 mM L-glutamine, 10% FBS, and 100 U penicillin/streptomycin.

### Preparation and assessment of polarized H292 monolayers

Polarized H292 monolayers were prepared as previously described (39). The transepithelial resistance of lung epithelial monolayers is typically very low (unpublished data). Hence, to assess the generation of intact, confluent H292 monolayers, for a sampling of polarized monolayers, we measured the ability of horseradish peroxidase (HRP) added to the basolateral compartment to be detected in the apical chamber after 20 minutes, as previously described (40). Potential defects in Transwell monolayer integrity were assessed after collagen coating via detecting excessive buffer loss and after cell seeding via changes in pH of cell media. Approximately 3% of H292 monolayers were excluded from our experiments.

### Toxin purification

The expression and purification of recombinant toxins and toxin derivatives from *Escherichia coli* were carried out as previously described (41) with the following modifications. Growth of *E. coli* XL-1 Blue containing pRT20 or derivatives thereof was initiated by inoculating 1 L of sterile TB broth with a 1:100 inoculum of an overnight culture grown at room temperature with 100 μg/mL ampicillin. The 1 L culture was incubated at 37 °C with shaking at 200 rpm. At OD_600_ 0.5-0.6, expression of toxin was induced by the addition of isopropyl β-D-thiogalactopyranoside (IPTG, Fisher Scientific) to a final concentration of 1 mM. Ampicillin was also added to a final concentration of 100 μg/mL. The induced culture was grown overnight at room temperature and pelleted by centrifugation. Cell pellets were resuspended in a total of 30 mL of PBS. Halt Protease Inhibitor Cocktail (Thermo Fisher Scientic) was added to prevent proteolytic degradation of the toxin. Cells were lysed by two passages through a microfluidizer, and cell debris was removed by centrifugation at 10,000 x *g* for 30 min at 4 °C. The supernatant containing the polyhistidine-tagged toxin was loaded onto a Ni-NTA agarose column (Qiagen). The column was then washed with a 20–120 mM gradient of imidazole to remove additional contaminating proteins. Bound toxin was then eluted (2 mL/min) with 10 mL of PBS containing 500 mM imidazole. SDS-PAGE was performed on the collected fractions to confirm toxin purity. 10% (v/v) glycerol and 5 μM Tris(2-carboxyethyl)phosphine hydrochloride (TCEP) was added to toxin-containing fractions, which were then flash-frozen and stored at −80 °C until use.

### Toxin activity and cell viability by FACS

Toxin aliquots were thawed on ice and then spun at 14,000 rpm in a microcentrifuge at 4 °C to remove protein precipitate for 10 min. Toxin concentration was determined using Bradford Reagent (BioRad) per the manufacturer’s instructions. 5 x 10^5^ H292 cells were placed in 100 µl volumes in a non-adherent 96 well plate and were treated for 15 minutes at 37°C with toxin at varying concentrations for activity determination, or with *S. pneumoniae* at MOI of 2 or 20 to assess the effect of bacterial infection on membrane permeability. Addition of 5% Triton-X 100 and HBSS+ were used as positive and negative lysis controls, respectively. For PMN viability assessment, 5 x 10^5^ human peripheral blood PMNs were plated per well instead. The plate was then spun at 1200rpm for 5 minutes and cells were resuspended in 1 mg/ml of propidium iodide (PI). Cells were filtered through 100 μM filters into 100 μl of PBS with 10% serum, placed on ice, and kept in the dark until analysis by flow cytometry. Cells were run through a FACSCalibur flow cytometer (BD Biosciences) and a minimum of 5 x 10^4^ events were analyzed per replicate. For determining toxin activity unit, collected data were analyzed using FlowJo software (Tree Star, Inc.) to determine the toxin concentration that resulted in 50% of the H292 cells becoming PI+. This concentration was defined as “1 Unit” of toxin activity.

### Infection and toxin treatment of polarized H292 monolayers

*S. pneumoniae* grown to log phase were washed and resuspended to indicated concentrations in Hanks’ balanced salt solution (HBSS) supplemented with Ca^2+^ and Mg^2+^. 25 μl of bacterial suspension was added to the apical surface of inverted Transwells and incubated at 37°C with 5% CO_2_ for 2.5 hours to allow for attachment and infection of monolayers. To assess for bacterial adhesion, monolayers were infected with GFP-expressing *S. pneumoniae* for 2.5 h hours then fixed in 4% PFA. Adhered bacteria were visualized by IF microscopy and bound bacteria per field of view was used to determine total bound bacteria per monolayer. For toxin treatments, 50 μl of the indicated units of toxin were added to the apical surface of inverted Transwells for 1 h and incubated at 37°C with 5% CO_2_. After treatment, Transwells were washed and placed in 24-well plates containing HBSS+Ca^2+^/ Mg^2+^ and incubated for an additional 2.5 h to allow for bacteria translocation. Permeability to horseradish peroxidase (HRP) was used to assess monolayer barrier integrity post treatment. Buffer in the basolateral chambers were sampled and bacterial translocation across the monolayer was evaluated by serial dilutions and plating on blood agar. Bacterial translocation index was calculated as total colony forming unites (CFUs) in basolateral chamber normalized to the initial inoculum.

### PMN transepithelial migration assay

1 x 10^6^ PMNs, isolated from whole blood obtained from healthy human volunteers, were added to the basolateral chamber after infection of the monolayer with *S. pneumoniae* and allowed to transmigrate for 2.5 hours. PMNs in the apical chamber were quantified by myeloperxidase (MPO) assay, as described (42). Briefly, monolayers and their underlying Transwell filters were removed from individual wells of a 24-well plates, leaving the 24-well plate containing the apical buffer and migrated neutrophils for each sample and control well. To assess viability of transmigrated PMNs in the apical chamber, 20 μl of apical chamber buffer was sampled and incubated with 3,3’,5,5’-Tetramethylbenzidine (TMB) substrate for detection of released MPO activity. Then, 50 μl of 10% Triton X-100 was added to the apical chamber and gently rocked for 20 min at 4°C to lyse all transmigrated PMNs. 50 μl of citrate buffer was added to each sample and the 24-well plate was gently rocked for 20 min at 4°C. ABTS solution was freshly prepared and 50 μl of hydrogen peroxide was added to the 2,2’-azino-bis(3-ethylbenzothiazoline-6-sulphonic acid) (ABTS) solution. 100 μl from each well was transferred to a 96-well plate and 100 μl of ABTS solution was added to each sample in the 96-well plate. The 96-well plate was incubated in the dark at room temperature for 5-10 min until it was read on a microplate reader for absorbance at a wavelength of 405 nm. Absorbance measurement was converted to neutrophil number using a standard curve and used to determine percent PMN transmigrated.

### Fluorescence microscopy assessment of monolayer viability and integrity

Monolayer viability and integrity were assessed by fluorescence microscopy of DAPI for cell confluency, E-cadherin for adherens junction organization, and PI for cell permeabilization. To prepare samples for fluorescence microscopy, monolayers were fixed in 4% PFA and stained with 1 mg/ml PI. Another set of fixed monolayers were permeabilized with 0.1% Triton-X 100 in PBS plus 3% BSA, and stained with DAPI, phalloidin, and anti-E-cadherin (24E10), followed by α-rabbit-FITC secondary antibody, and visualized on excised filters. To visualize cell surface exposed E-cadherin, monolayers were stained with ectodomain E-cadherin antibody (DCMA-1) followed by α-rat-FITC secondary antibody, and then permeabilized with 0.1% Triton-X 100 in PBS plus 3% BSA and stained with DAPI (blue) and phalloidin (red). Quantification of E-cadherin junction organization was carried out with the python script IJOQ as previously described (43). Briefly, the script quantifies spatial distribution of a junctional molecule along the continuous contacts with neighboring cells by determining the degree of intersection between networks of the molecule to a predefined grid-like network. Percentage epithelial cell loss was quantitated by count of DAPI stained epithelial cell nuclei per field of view.

### Western blot

Transwell monolayers were washed three times in HBSS, excised, and lysed in 100 μl of Pierce radioimmunoprecipitation assay (RIPA) buffer (Thermo Fisher Scientific) containing protease inhibitors (cOmplete Mini; Roche Diagnostics, Indianapolis, IN). Proteins were resolved by SDS-PAGE under reduced conditions and transferred onto a polyvinylidene difluoride (PVDF) membrane for immunoblotting. Membranes were blocked in Tris-buffered saline–1% Tween (TBS-T) with 5% nonfat milk, and subsequently incubated with the E-cadherin antibody (24E10) at 1:1000. Following washing, immunoblots were probed with the appropriate secondary α-rabbit antibody conjugated to horseradish peroxidase at a dilution of 1:5000. Immune complexes were detected using the SuperSignal West Pico Chemiluminescent Substrate (ThermoFisher Scientific), following the manufacturer’s instructions, and imaged using a Syngene G:Box XR5 imager. Densitometry analysis was carried out with ImageJ FIJI, normalized to the respective beta actin loading control.

### Matrix metalloprotease (MMP) activity

PLY-treated H292 monolayers or PMNs in suspension were treated with 1U PLY and incubated for 30 min at 37 °C with 10 μM fluorogenic peptide MMP substrate (Mca-PLAQAV-Dpa-RSSSR-NH2, R&D Systems) diluted in 25 mM Tris buffer, pH 8.0. Fluorescence intensity over time was read on a BioTek Synergy HT plate reader and area under the curve of kinetic substrate conversion was used to determine MMP activity.

### Ethics statement

All work with human donors was approved by the Tufts University Human Investigation Review Board (IRB). Written informed consent was obtained from all donors and collection was carried out with the assistance of services provided by the Tufts Clinical and Translational Research Center (CTRC).

### Presentation of data and statistical analyses

Due to intrinsic donor-to-donor variability of human PMNs and their transmigration, efficiency of transmigration was compared within individual experiments but not between experiments. The conclusions drawn were those found to be reproducible and statistically significant across independent experiments. Statistical analysis was carried out using the program GraphPad Prism (GraphPad Software, San Diego, CA). p values <0.05 were considered significant in all cases. For all graphs, the mean values +/− SEM are shown.

## Results

### High dose *S. pneumoniae* infection elicits PLY-dependent PMN recruitment, which promotes epithelial detachment and bacterial translocation

Previously, we reported that PMN transmigration, is required for retrograde translocation of *S. pneumoniae* across polarized airway epithelial monolayers *in vitro* and for bacteremia in mice (21, 31). We further showed that PMN transmigration is promoted by PLY-induced secretion of the inflammatory eicosanoid HXA_3_ (20). Here, we utilized polarized H292 cells grown on 5 µm pore size Transwell filters to directly test whether PLY-promoted PMN transmigration promotes barrier disruption and subsequent bacterial translocation. We first infected monolayers apically with TIGR4 at a calculated multiplicity of infection (MOI) of 20 using 1 x 10^7^ total bacteria per Transwell (See Materials and Methods), an infectious dose used in previous reports (20, 21). This resulted in 5 x 10^5^ total bound bacteria per Transwell with indistinguishable binding efficiency between wild type (WT) TIGR4 and PLY deficient TIGR4*Δply*, as visualized by immunofluorescence (IF) microscopy (Supplemental Figure 1A). Infection with WT TIGR4 triggered the basolateral-to-apical transmigration of ∼30 x 10^4^ human PMNs, accounting for 30% of PMNs added to the basolateral chamber (Figure 1A, “WT”). This induction of PMN transmigration was partially PLY-dependent, as the TIGR4*Δply* strain was associated with a ∼1.5-fold reduction in PMN movement compared to the WT strain (p<0.01) (Figure 1A, “*Δply*”). Nevertheless, the level of TIGR4*Δply* induced PMN transmigration was 14-fold (p<0.0001) above the mock-infected background level, confirming that PMN transmigration can be elicited in PLY-dependent and -independent manners (21, 31).

**Figure 1.**
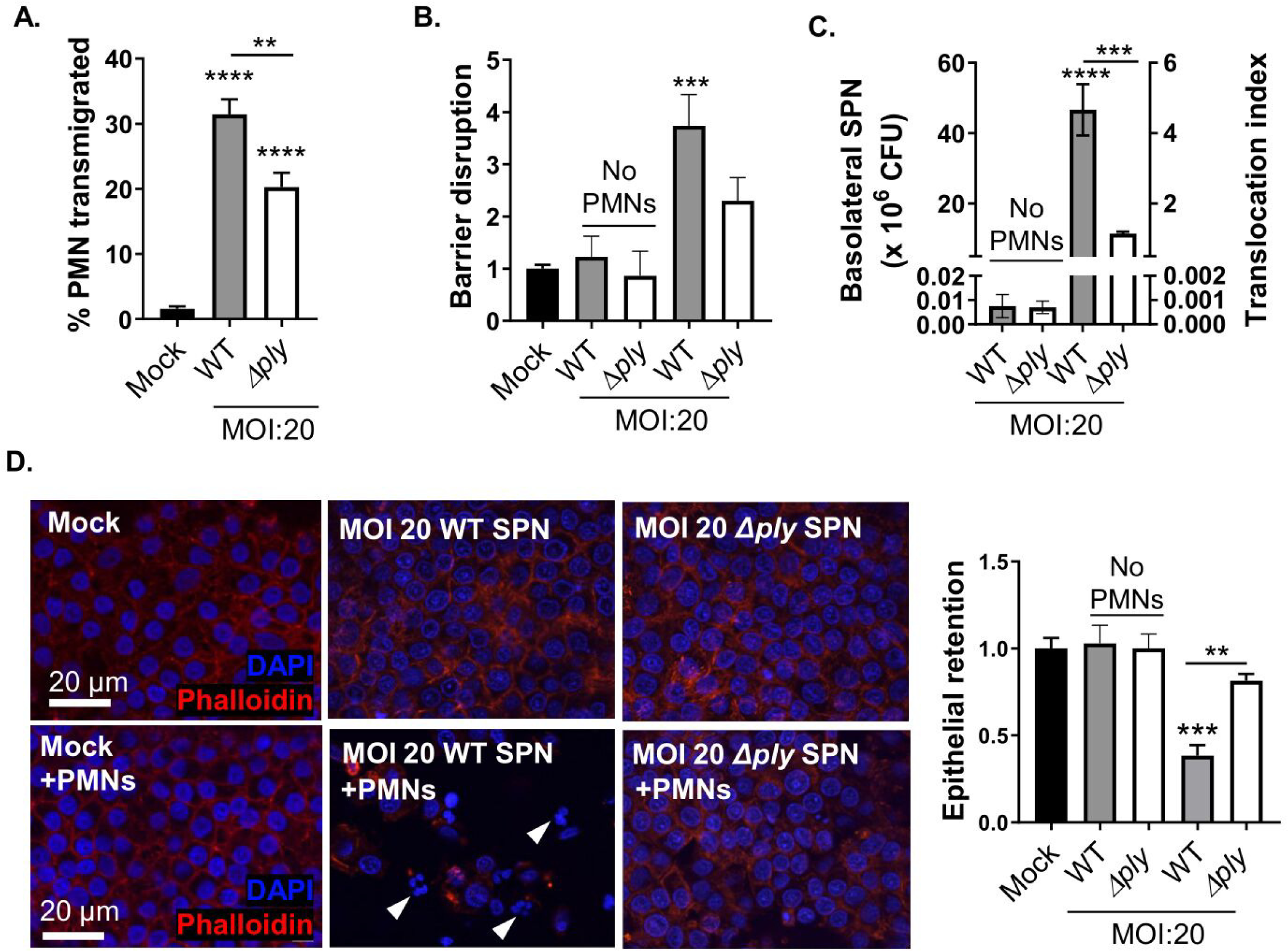
High dose *S. pneumoniae* infection elicits PLY-dependent PMN recruitment, which promotes epithelial detachment and bacterial translocation. **A.** 1 x 10^6^ human PMNs were added to the basolateral side of Transwell seeded polarized H292 monolayers that were mock-infected (with HBSS, “Mock”), infected apically with wild type TIGR4 (“WT”), or infected apically with a pneumolysin-deficient mutant (“*Δply*”) at the indicated MOI for 2.5 h. Subsequently, PMN transepithelial migration after 2.5 h was quantified by MPO activity. **B.** Barrier integrity of monolayers was measured by HRP flux post-infection, normalized to mock-infected. **C.** Transepithelial migration of bacteria was quantitated by plating for CFUs, and the translocation index was determined by normalizing SPN present in the basolateral compartment to the infection inoculum. **D.** Fluorescence microscopy of monolayers stained with DAPI (blue) and phalloidin (red). White arrows indicate PMN nuclei. Scale bar = 20 μm. Epithelial cell retention was quantified by epithelial cell nuclei number per field of view. Each panel shown is representative of three independent experiments in triplicates, or pooled data from at least three independent experiments. Error bars represent mean ± SEM. Statistical analysis was performed using ordinary one-way ANOVA: \**p-value* ≤ 0.05, \*\**p-value*≤ 0.01, \*\*\**p-value* ≤ 0.001, \*\*\*\**p-value* ≤ 0.0001.

To examine the influence of PLY-dependent and -independent PMN transmigration on barrier dysfunction, we measured the apical-to-basolateral flux of horseradish peroxidase (HRP) and the apical-to-basolateral movement of bacteria. Infection by WT TIGR4 in the presence of PMNs was associated with a three-fold increase in HRP flux (p<0.001) (Figure 1B, “WT”) and the detection of 4.8 x 10^6^ *S. pneumoniae* in the basolateral chamber by quantitation of colony-forming units (CFUs) (p<0.0001) (Figure 1C, “WT”), which was approximately 7000-fold more than the number of translocated bacteria in the absence of PMNs (Figure 1C, “No PMNs”, “WT”). The WT TIGR4 translocation index, defined as the number of bacteria in the basolateral chamber at five hours (2.5 h infection followed by 2.5 h of PMN co-incubation) post-infection divided by the initial inoculum, was 4.8 in the presence of PMNs, as opposed to ∼0.0007 in the absence of PMNs (Figure 1C), indicating efficient translocation only in the presence of PMNs, consistent with our previous findings (21, 31). (The calculated translocation index of greater than 1 indicates that there is some degree of bacterial replication during the five-hour incubation). Infection with TIGR4*Δply*, which elicited ∼1.5-fold less PMN migration than WT TIGR4 (Figure 1A), led to an equivalently lower level of HRP flux, but this difference did not reach statistical significance (Figure 1B). The degree of HRP flux induced by TIGR4*Δply* was two-fold greater than baseline HRP flux in mock-infected monolayers, but this difference also did not reach statistical significance (Figure 1B). Barrier disruption by PMNs in response to TIGR4*Δply* infection was most clearly revealed by bacterial translocation, because 10 x 10^6^ TIGR4*Δply* were present in the basolateral chamber, resulting in a translocation index of 1, and an approximately 1000-fold increase in bacterial transmigration in comparison to that in the absence of PMNs (Figure 1C). Thus, PMN transmigration, whether induced in a PLY-dependent or PLY-independent manner, promotes *S. pneumoniae* transmigration.

To visualize how PMN transmigration to *S. pneumoniae* disrupted the epithelial barrier, we assessed monolayer confluency after *S. pneumoniae* infection and PMN transmigration by immunofluorescence (IF) staining of host cell nuclei (DAPI) and F-actin (phalloidin). At a MOI of 20, infection by WT TIGR4 or TIGR4*Δply* did not alter monolayer confluency (Figure 1D, top row), but did result in approximately 50% of cells becoming permeable to propidium iodide (PI) upon WT TIGR4 infection (Supplemental Figure S1B). TIGR4*Δply* infection, on the other hand, did not result in significant PI staining, suggesting that PLY permeabilized cells during infection (Supplemental Figure S1C). Although we and others have shown that H292 cells initiate membrane repair processes to prevent cell death following PLY insult (20, 44)), it is possible that PLY-permeabilized cells are particularly vulnerable to dislocation from the monolayer upon subsequent addition of PMNs. We thus examined monolayer confluency by IF again post PMN transmigration. PMN infiltration to WT TIGR4 infected monolayers can be identified by their multi-lobed nuclei (Figure 1D, white arrowheads), and PMN migration was associated with loss of 62% of epithelial cells from the monolayer (Figure 1D, bottom row, “Mock” vs. “WT SPN”, with image analysis quantitation on right). In contrast, infection by TIGR4*Δply* resulted in a 20% loss of epithelial cells, significantly less than the loss associated with infection by WT TIGR4 (p<0.01) (Figure 1D).

Because PLY can also permeabilize PMNs, we used flow cytometry to evaluate the degree of PI staining after direct infection of PMN with WT or TIGR4*Δply* and observed a PLY-dependent permeabilization of 2% of PMNs (Supplemental Figure 1D). We also evaluated PMN permeabilization after traversing the monolayer by measuring MPO activity in the apical buffer immediately after PMN transmigration. We observed a 2-fold and significant increase in MPO activity in WT TIGR4 infected Transwells (Supplemental Figure 1E), which corresponds to if approximately 10% of transmigrated PMNs released MPO. No MPO activity above background was observed with TIGR4*Δply* infection. This raises the possibility that at an MOI of 20, PMN release of effector molecules post transmigration may contribute to epithelial detachment. These findings implicate PMN transmigration, augmented by PLY, as a key factor that drives monolayer disruption, barrier integrity compromise, and *S. pneumoniae* translocation.

### Low dose infection elicits PLY-dependent PMN recruitment, which drives *S. pneumoniae* translocation in the absence of epithelial detachment

PMN transmigration in response to TIGR4 infection is highly MOI-dependent (21, 31). Hence, we performed a similar series of experiments infecting H292 monolayers with a 10-fold lower infection dose, which resulted in approximately 1 x10^5^ total bacteria bound/Transwell (Supplemental Figure 1A). WT TIGR4 infection at MOI 2 triggered transmigration of ∼15 x 10^4^ human PMNs (i.e., 15% of total PMNs added; Figure 2A, “WT”), approximately two-fold lower than that observed at a MOI of 20 (Figure 1A, “WT”), yet 30-fold higher (p<0.01) compared to mock-infected monolayers. Infection with TIGR4*Δply* resulted in an approximate six-fold (and not statistically significant) increase in PMN transmigration compared to mock-infected monolayers, indicating that PMN transmigration remains PLY-promoted at the lower MOI (Figure 2A, “*Δply*”). Notably, this PLY-dependent effect was not due to more efficient bacteria-host cell interaction because monolayer binding by TIGR4*Δply* was indistinguishable from wild type (Supplemental Figure 1A), nor due to host cell killing by PLY because epithelial and neutrophil viability was not affected at this MOI (Supplemental Figure 1B-E). Disruption of barrier integrity correlated with PMN transmigration, because HRP flux was elevated 2.2-fold (p<0.01) upon infection with WT TIGR4 but not significantly by TIGR4*Δply* compared to mock-infected monolayers (Figure 2B). Correspondingly, at MOI 2, 0.9 x 10^6^ WT TIGR4 and 0.2 x 10^6^ TIGR4*Δply* bacteria migrated to the basolateral chamber respectively (Figure 2C). While the lower number of bacteria translocated was expected given the lower inoculum, it is notable that translocation was still PMN-promoted, because in the absence of PMNs, translocation of WT TIGR4 and TIGR4*Δply* was diminished 700- and 300-fold, respectively (Figure 2C). Monolayer confluency was retained despite robust PMN and *S. pneumoniae* transmigration (Figure 2D). Hence, at a MOI of 2, PLY and PMNs promoted bacterial migration in a manner that did not require overt destruction of the monolayer.

**Figure 2.**
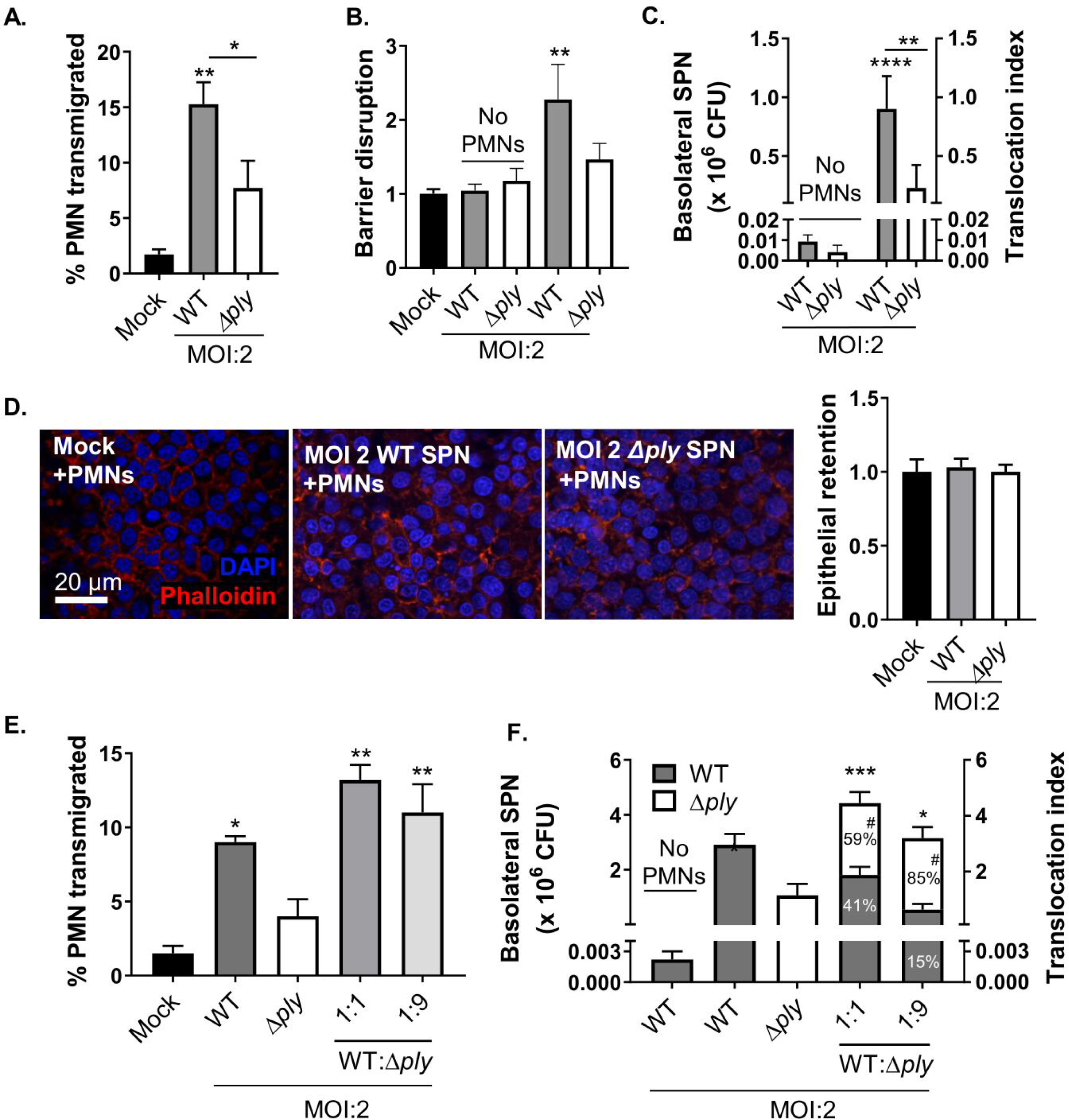
Low dose infection elicits PLY-dependent PMN recruitment, which drives *S. pneumoniae* translocation in the absence of epithelial detachment. **A.** 1 x 10^6^ human PMNs were added to the basolateral side of monolayer mock-infected (with HBSS “Mock”) or infected apically with WT or *Δply* SPN at indicated MOI. PMN transepithelial migration was quantified by MPO activity. **B.** Barrier integrity of monolayers was measured by HRP flux post-infection, normalized to mock-infected. **C.** Transepithelial migration of bacteria was quantitated by plating for CFUs, and translocation index was determined by normalizing SPN present in basolateral chamber to infection inoculum. **D.** Fluorescence microscopy of monolayers stained with DAPI (blue) and phalloidin (red). Scale bar = 20 μm. Epithelial cell retention was quantified by epithelial cell nuclei number per field of view. **E.** Monolayers were mock-infected (HBSS) or apically infected apically with WT, *Δply*, a 1:1 mix of WT and *Δply*, or a 1:9 mix of WT and *Δply* SPN at indicated MOI. PMN transepithelial migration was quantified by MPO activity. **F.** Transepithelial migration of bacteria was quantitated by plating, and translocation index was determined by normalizing SPN migration to infection inoculum. Statistical analysis comparing total SPN migration between respective groups indicated with ‘*’, statistical analysis comparing TIGR4*Δply* migration between respective groups indicated with ‘#’. Each panel shown is representative of three independent experiments in triplicates, or pooled data from at least three independent experiments. Error bars represent mean ± SEM. Statistical analysis was performed using ordinary one-way ANOVA: \**p-value* ≤ 0.05, \*\**p-value*≤ 0.01, \*\*\**p-value* ≤ 0.001.

To determine whether the pro-transmigration functions of PLY were restricted to PLY-producing bacteria or could function in *trans*, we infected H292 monolayers with either a 1:1 or 1:9 mixture of WT TIGR4 and TIGR4*Δply*, where TIGR4*Δply* can be identified through chloramphenicol resistance. To avoid the epithelial detachment that is associated with the MOI 20 dose (Figure 1D), we utilized the MOI of 2 dose. As shown previously in Figure 2A, WT TIGR4 elicited a stronger PMN response than TIGR4*Δply* (Figure 2E). Both the 1:1 and the 1:9 mixed infections led to neutrophil recruitment similar to that of WT TIGR4-infected monolayers (Figure 2E), consistent with previous reports of PLY-mediated PMN chemoattractant HXA_3_ secretion by host cells (20). In the presence of transmigrating PMNs, bacteria from both the 1:1 and 1:9 mixed infections efficiently migrated to the basolateral compartment with efficiencies similar to that of infections with WT TIGR4 only (Figure 2F). Differential antibiotic resistance permitted distinction between the two strains and revealed that the transmigration of TIGR4*Δply* was significantly (p<0.05, indicated by ‘#’) increased in the presence of WT TIGR4 (Figure 2F). Moreover, the ratio of WT:*Δply* bacteria in the basolateral chamber was similar to input ratios upon apical infection, i.e., the 1:1 WT:*Δply* mixture resulted in a 1 : 1.4 output and the 1:9 input resulted in a 1 : 5.7 output (Figure 2F). Thus, the pro-translocation activity of PLY promotes migration of both PLY-producing and PLY-deficient bacteria, likely by eliciting epithelial cell HXA_3_ production and concomitant PMN migration, which contributes to barrier disruption(20).

### *S. pneumoniae* infection results in cleavage and mislocalization organization of the AJ protein E-cadherin via PLY-dependent and PLY-independent pathways

*S. pneumoniae* have been reported to target endothelial and epithelial junctional components to compromise respiratory barrier function (4). To determine if *S. pneumoniae* targets junctional components in our model, we infected H292 cell monolayers and stained with an antibody that recognizes the extracellular domain of E-cadherin, a transmembrane AJ protein capable of homotypic binding to promote barrier formation that is robustly expressed by H292 cells. When visualized by IF microscopy, E-cadherin is located in circumferential rings at the cell periphery (Figure 3A, “Mock”; supplemental Figure 2). Addition of PMNs to a mock-infected monolayer had no effect on E-cadherin localization of staining intensity visualized by IF (Figure 3A, “Mock+PMNs”).

**Figure 3.**
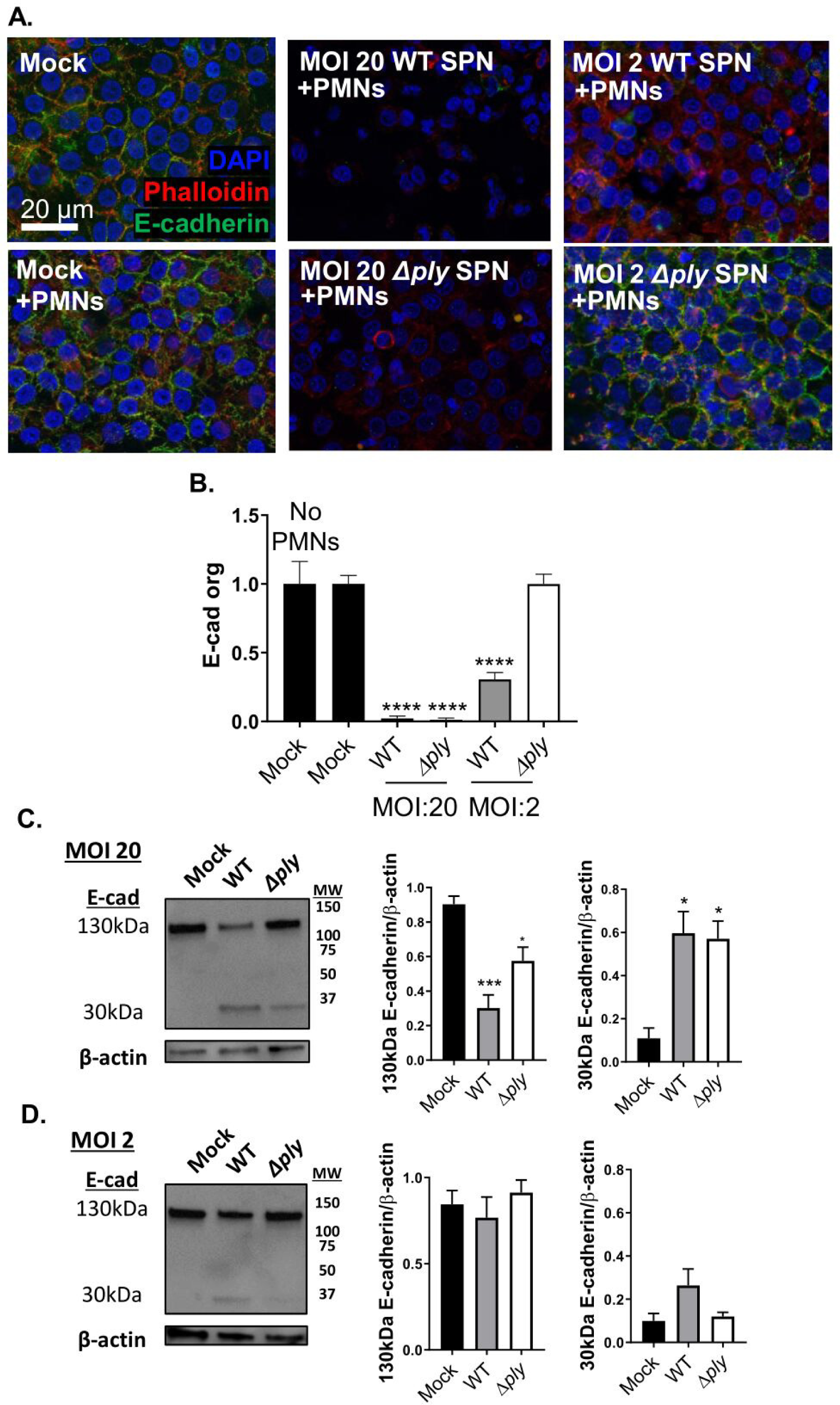
*S. pneumoniae* cleaves and disrupts the localization of E-cadherin via PLY-dependent and PLY-independent pathways. **A.** 1 x 10^6^ human PMNs were added to the basolateral side of monolayers mock-infected (HBSS) or infected apically with WT or *Δply* SPN at indicated MOI. Monolayers were stained with ectodomain E-cadherin antibody (green) before permeabilization and with DAPI (blue) and phalloidin (red) after permeabilization. Scale bar = 20 μm. **B.** Peripheral E-cadherin organization was quantitated by image analysis with intercellular junction organization quantification (IJOQ). Each panel shown is representative of three independent experiments in triplicates. **C.** MOI 20 and **D.** MOI 2 infected monolayers were lysed for western blotting of E-cadherin and desitometry analysis normalized to β-actin. Error bars represent mean ± SEM. Statistical analysis was performed using ordinary one-way ANOVA: \*\*\*\**p-value* ≤ 0.0001.

To examine whether H292 cell E-cadherin complexes are disrupted upon PMN migration induced by *S. pneumoniae*, monolayers were apically infected with WT TIGR4 or TIGR4*Δply* at an MOI of 20 or 2. WT TIGR4 infected monolayers lost all cell periphery E-cadherin, regardless of infection dose (Figure 3A, top row, “WT SPN”; supplemental Figure 2). In contrast, TIGR4*Δply* was able to induce loss pericellular E-cadherin localization at a MOI of 20 but not at a MOI of 2 (Figure 3A, bottom row, “*Δply* SPN”). Quantification of E-cadherin localization specifically at the cell periphery was carried out with the Intercellular Junction Organization Quantification (IJOQ) script (43), which quantifies the continuity in pericellular staining. This analysis showed both WT TIGR4 and TIGR4*Δply* infection at a MOI of 20 eliminated E-cadherin organization (p<0.0001), whereas at a MOI of 2, only WT TIGR4 infection induced a significant (60%) reduction in E-cadherin organization (p<0.0001) (Figure 3B).

To determine if E-cadherin structure is altered by infection by *S. pneumoniae*, we performed western blotting of E-cadherin in total cell lysates from uninfected or infected monolayers. This revealed a PLY-dependent decrease in the 130 kDa full length E-cadherin and the appearance of an E-cadherin-related 30 kDa species after infection with WT TIGR4 at MOI 20, (Figure 3C). At MOI 2, infection by WT TIGR4 also resulted in apparent E-cadherin cleavage, although densitometric analysis did not reveal statistical significance (Fig. 3D). E-cadherin cleavage appeared to be partially dependent on PLY, because infection by TIGR4 *Δply* at either MOI 2 or 20 resulted in less cleavage than during WT infection (Figs. 3C and D, “*Δply*”). Thus, E-cadherin cleavage and mislocalization is triggered via both PLY-dependent and PLY-independent pathways, with the PLY-dependent pathway capable of triggering these changes at a lower infectious dose.

### PMNs are not required for E-cadherin mislocalization by *S. pneumoniae* at either low or high infectious dose

Independent of PMNs, *S. pneumoniae* has been reported to downregulate epithelial junctional components during infection through mechanisms such as TLR2 engagement (9) and pneumococcal neuraminidase activity (15). To determine whether PMNs are required for the PLY-dependent and - independent E-cadherin mislocalization described above, we infected monolayers with WT TIGR4 and TIGR4*Δply* at a MOI of 20 or 2 in the absence of PMNs. At a MOI of 20, both WT TIGR4 and TIGR4*Δply* depleted all pericellular E-cadherin (Figure 4A), just as when PMNs were present (Figure 3A). Lowering the MOI to 2, we observed that WT TIGR4 induced a 38% (p < 0.01) decrease in E-cadherin localization to cell periphery (Figure 4B, top row, “MOI 2 WT SPN”, and right, solid bars). In contrast, TIGR4*Δply* infection at a MOI of 2 had no effect on E-cadherin localization (Figure 4B, top row, “MOI 2 *Δply* SPN”, and right, solid bars).

**Figure 4.**
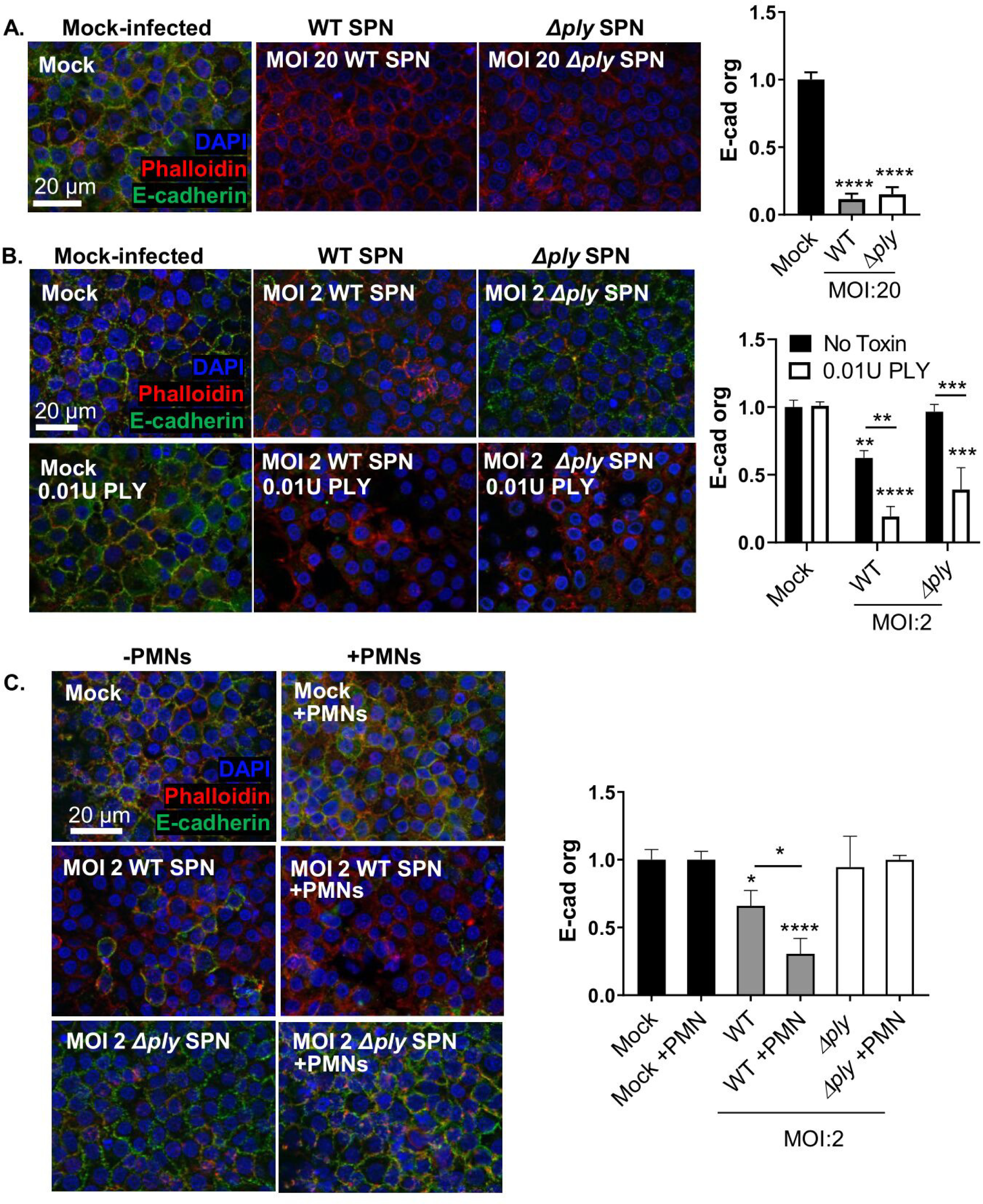
PMNs are not required for E-cadherin mislocalization by *S. pneumoniae* at either low or high infectious dose. **A.** Fluorescence microscopy of monolayers mock-infected (HBSS) or infected apically with WT or *Δply* SPN at indicated MOI in the absence of PMNs, stained with ectodomain E-cadherin antibody (green) before permeabilization and with DAPI (blue) and phalloidin (red) after permeabilization. Scale bar = 20 μm. Peripheral E-cadherin organization was quantitated by image analysis with IJOQ. **B.** Fluorescence microscopy of monolayers mock-infected (HBSS) or infected apically with WT or *Δply* SPN at indicated MOI, and complemented with recombinant PLY where indicated. Monolayers were stained with ectodomain E-cadherin antibody (green) before permeabilization and with DAPI (blue) and phalloidin (red) after permeabilization. Scale bar = 20 μm. Peripheral E-cadherin organization was quantitated by image analysis with IJOQ. **C.** Fluorescence microscopy of monolayers mock-infected with HBSS (mock) or infected apically with WT or *Δply* SPN at indicated MOI in the presence or absence of PMNs, stained with ectodomain E-cadherin antibody (green) before permeabilization and with DAPI (blue) and phalloidin (red) after permeabilization. Scale bar = 20 μm. Peripheral E-cadherin organization was quantitated by image analysis with IJOQ. Each panel shown is representative of three independent experiments in triplicates. Error bars represent mean ± SEM. Statistical analysis was performed using ordinary one-way ANOVA: \**p-value* ≤ 0.05, \*\**p-value*≤ 0.01, \*\*\**p-value* ≤ 0.001.

To determine if *S. pneumoniae* disruption of E-cadherin integrity could be augmented by the addition of purified PLY, we added recombinant PLY at a sublytic PLY concentration of 0.01 toxin activity units (Materials and Methods) (20), a concentration that by itself was incapable of disrupting E-cadherin localization (Figure 4B, “Mock, 0.01U PLY”, and right, open bars). We found that, when combined with infection by WT TIGR4, this level of exogenous PLY enhanced E-cadherin disruption from 38% to 81% (p < 0.01; Figure 4B, bottom row, “WT SPN, 0.01U PLY”, and right, open bars). Whereas TIGR4*Δply* at a MOI of 2 was insufficient to perturb E-cadherin on its own, the addition of 0.01U PLY to monolayers infected with TIGR4*Δply* resulted in a 53% (p < 0.001) reduction in E-cadherin cell periphery organization (Figure 4B, “Mock+0.01U PLY”, and right, open bar).

Because PLY promotes PMN transmigration (18, 20), we hypothesized that the addition of PMNs to monolayers infected with WT TIGR4 may further compromise E-cadherin integrity. Indeed, the addition of PMNs to the basolateral chamber of monolayers infected with WT TIGR4 at a MOI of 2 resulted in a two-fold decrease (p<0.05) in E-cadherin organization (Figure 4C, “WT” vs. “WT + PMN”). As expected, PLY was critical to this augmentation, because the addition of PMNs to monolayers infected with TIGR4*Δply* had no effect (Figure 4C, “*Δply*” vs. “*Δply* + PMN”).

### Pore formation by PLY and related toxins is necessary and sufficient for E-cadherin mislocalization

To investigate whether purified PLY is sufficient for disrupting E-cadherin localization, we examined E-cadherin organization after apical treatment of H292 monolayers with various concentrations of PLY. 1U of PLY, a concentration resulting in transient permeabilization of 50% of host cell followed by membrane repair (20), ablated pericellular E-cadherin (Figure 5A). A 10-fold lower PLY concentration also significantly (p < 0.05) reduced E-cadherin organization compared to mock treatment (Figure 5A). 0.01U of PLY had no effect on E-cadherin organization (Figure 5A). Thus, purified PLY mimics E-cadherin disruption by live *S. pneumoniae*, and in an infection dose-dependent manner. Western blotting of E-cadherin confirmed no cleavage of E-cadherin with 0.01U PLY treatment, partial diminution of full-length E-cadherin and the appearance of a very faint 30 kDa species with 0.1U PLY treatment, and a dramatic reduction in full length E-cadherin and the appearance of the 30kDa cleavage product with 1U PLY treatment (Figure 5B).

**Figure 5.**
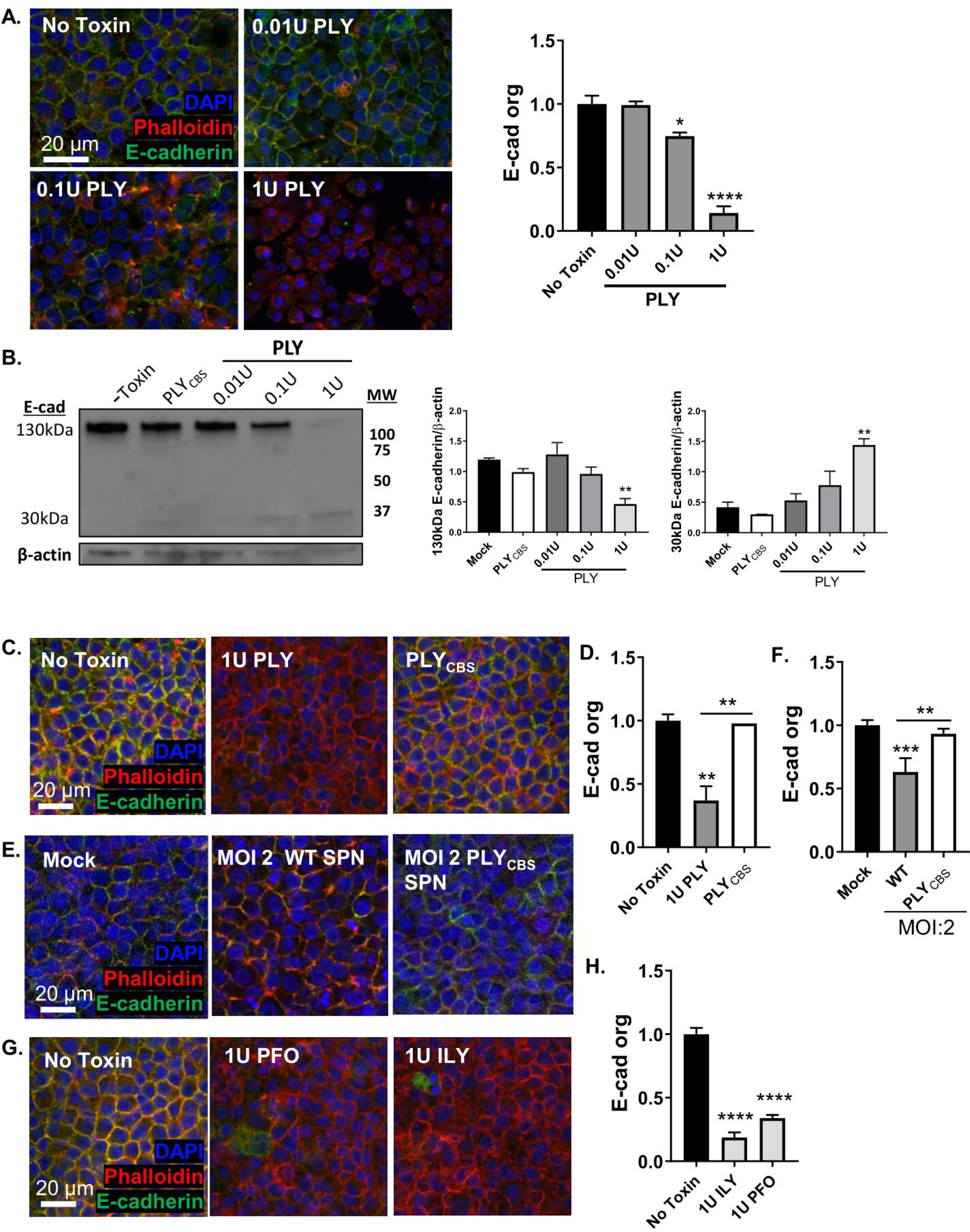
Treatment of epithelial cells by PLY or related pore-forming toxins is sufficient to induce cleavage and mislocalization of E-cadherin. Monolayers were treated with HBSS (No Toxin) or the indicated units of recombinant PLY.**A.** Monolayers were stained with ectodomain E-cadherin antibody (green) before permeabilization and with DAPI (blue) and phalloidin (red) after permeabilization. Scale bar = 20 μm. Peripheral E-cadherin organization was quantitated by image analysis with IJOQ. **B.** Monolayers were lysed for western blotting of E-cadherin and desitometry analysis normalized to β-actin. **C.** Fluorescence microscopy of monolayers treated apically with purified PLY_CBS_ protein at equimolar concentrations to 1U of PLY. Monolayers were permeabilized and stained with E-cadherin antibody (green), DAPI (blue), and phalloidin (red). Scale bar = 20 μm. **D.** Peripheral E-cadherin organization was quantitated by image analysis with IJOQ. **E.** Fluorescence microscopy of monolayers infected apically with PLY_CBS_-expressing SPN (PLY_CBS_ SPN) at indicated MOI. Monolayers were permeabilized and stained with E-cadherin antibody (green), DAPI (blue), and phalloidin (red). Scale bar = 20 μm. **F.** Peripheral E-cadherin organization was quantitated by image analysis with IJOQ. **G.** Fluorescence microscopy of monolayers treated with the indicated units of purified PLY, intermedilysin (ILY) of *S. intermedius*, or perfringolysin O (PFO) of *C. perfringens*. Monolayers were permeabilized and stained with E-cadherin antibody (green), DAPI (blue), and phalloidin (red). Scale bar = 20 μm. **H.** Peripheral E-cadherin organization was quantitated by image analysis with IJOQ. Each panel shown is representative of three independent experiments in triplicates. Error bars represent mean ± SEM. Statistical analysis performed using ordinary one-way ANOVA: \**p-value* ≤ 0.05, \*\**p-value* ≤ 0.01, \*\*\**p-value* ≤ 0.001, \*\*\*\**p-value* ≤ 0.0001.

Matrix metalloproteases (MMPs), which can be released by epithelial cells or PMNs during infection, are capable of cleaving extracellular matrix and junctional proteins, including E-cadherin (27, 45, 46). The pore-forming *Staphylococcus aureus* α-toxin (47, 48) and *Pseudomonas aeruginosa* exolysin (49) trigger E-cadherin cleavage in infected cells by activating host matrix metalloproteases (MMPs). PLY also has been shown to trigger MMP activation in A549 cells (47). We found that treatment of either epithelial cells or PMNs with 1 U of purified PLY triggered an increase of MMP activity in cell supernatants (Supplemental Figure 3A-B). This activity was inhibited by the broad spectrum MMP inhibitor GM6001. However, the MMP inhibitor GM6001 did not prevent E-cadherin cleavage (Figure 3C “GM6001”) or disruption of E-cadherin organization when monolayers were treated with 1U PLY or infected with WT TIGR4 at MOI 2 (Supplemental Figure 3C-D), implying that E-cadherin disuption observed was not mediated by MMPs.

Many of the biologic activities of PLY are mediated through its pore-forming activity (35), but PLY is also capable of altering cell behavior independent of pore-formation (34, 36). The mutant PLY^T459G•L460G^ (which we herein term “PLY_CBS_”) is mutated at two critical residues in the cholesterol binding site in PLY domain 4 and incapable of pore-formation (20, 50). H292 monolayers treated with PLY_CBS_ at equimolar concentration to 1U PLY had no effect on E-cadherin integrity by microscopy (Figure 5C-D). Likewise, apical infection with TIGR4 mutant producing PLY_CBS_ (PLY_CBS_ SPN), which is incapable of H292 cell permeabilization, failed to disrupt H292 E-cadherin organization (Figure 5E-F), indicating that pore-formation by PLY is essential for E-cadherin disruption.

PLY belongs to a family of cholesterol-dependent cytolysins (CDCs) that share substantial structural and functional conservation. We investigated whether other CDCs could also induce E-cadherin disruption. H292 monolayers apically treated with 1U of perfringolysin O (PFO) or intermedilysin (ILY), resulted in almost complete ablation of E-cadherin staining (Figure 5G-H), like that observed with 1U of PLY. These results indicate that the ability to disrupt pericellular localization of the epithelial cell AJ protein E-cadherin is a property conserved among CDCs and may be utilized by many CDC-expressing bacterial pathogens to target host cell junction networks.

## Discussion

The mucosal epithelium, sealed by a well-organized system of intercellular junctional complexes, is a physical line of defense against microbial invasion. Pathogens that cross epithelial barriers often disrupt these epithelial junctions, which facilitates tissue invasion by paracellular movement (51). *S. pneumoniae* has been shown to target airway junction proteins and compromise airway epithelial barrier function (4, 9, 10). Pneumococcal infection reduces alveolar occludin, ZO-1, claudin-5, and VE-cadherin in human lung explants (4), and downregulates claudin-7 and -10 in murine models (9). The reduction in critical junctional molecules can increase epithelial permeability and is likely an important step in facilitating pneumococcal dissemination (52); mice deficient in type I interferon, which show junctional defects in response to bacterial infection, are highly susceptible to *S. pneumoniae* bacteremia. (10). We thus examined the epithelial barrier and junctional integrity during *S. pneumoniae* infection of polarized H292 monolayers to systematically reveal requirements for *S. pneumoniae* translocation across the respiratory epithelium. We show that PLY and transmigrating PMNs 1) individually and cooperatively perturb localization and integrity of the AJ protein E-cadherin, 2) cause distinct degrees of epithelial damage depending on infection dose, PLY-mediated pore formation, and PMN presence, and 3) only when combined promote maximal bacterial cross-epithelial dissemination.

PLY, a *S. pneumoniae* virulence factor present in almost all clinical isolates of the species, increases alveolar permeability and alveolar epithelial cell injury (53). In human adenoid epithelial explant cultures, PLY ablates tight junctions and exposes host receptors for bacterial attachment (13). The notable invasiveness of serotype 1 pneumococci has been attributed to its rapid and robust release of PLY, which promotes ZO-1 disruption (14). Here, we found that *S. pneumoniae* at an MOI of 2 triggers both cleavage and mislocalization of the junctional protein E-cadherin in a manner dependent on PLY-mediated pore forming activity. Two other cholesterol-dependent cytolysins, ILY and PFO, induce intercellular junction disruption, suggesting that junction disruption is a conserved function of CDCs (54, 55). Other pathogens, such as *Staphylococcus aureus, Pseudomonas aeruginosa,* and *Serratia marcescens,* produce non-CDC pore-forming toxins that disrupt junctions *in vitro* (47, 49).

The ability of *S. pneumoniae* at an MOI (of 2) that did not compromise host cell membrane integrity indicates that E-cadherin disorganization is not a simple consequence of host cell death (20). PLY, like other pore-forming toxins, triggers membrane repair (44, 56–58), and, by activating GTPases that regulate actin assembly (11), may destabilize actin-associated junctional protein complexes (59, 60). PLY, although not a protease itself, has been shown to increase metalloprotease activity in A549 cells (47). Pore-forming toxins of *P. aeruginosa* and *S. marcescens* induce an influx of calcium ions, thus activating the calmodulin-regulated metalloproteinase ADAM-10, which cleaves E-cadherin (61). *S. aureus* toxin Hla directly binds and activates ADAM-10. Western blotting revealed a 30kDa cleavage product was generated upon *S. pneumoniae* infection or PLY treatment, suggesting disruption of E-cadherin organization is associated with its proteolytic cleavage. Consistent with previous reports (47, 48), we found that infection of PMNs or epithelial cells by PLY-producing SPN or treatment with purified PLY was associated with an increase in MMP activity in culture supernatants. However, inhibition of MMP did not prevent E-cadherin cleavage or mislocalization. Identification of proteases responsible for PLY-dependent cleavage of E-cadherin organization warrants future studies.

At an MOI of 20, both PLY-producing and PLY-deficient *S. pneumoniae* disrupted junctional organization of H292 cell monolayers. This PLY-independent pathway augments PLY-mediated disruption, as a PLY concentration (0.01U) that was insufficient for E-cadherin loss in isolation, in combination with PLY-deficient *S. pneumoniae* facilitated complete junctional disruption. TLR stimulants (9) and pneumococcal neuraminidase (15), which downregulate junctional components, are potential non-PLY bacterial factors that contribute to E-cadherin loss.

*S. pneumoniae*-mediated junctional disruption of monolayers alone was insufficient to promote bacterial translocation. Neutrophils secrete proteolytic enzymes capable of cleaving extracellular matrix and junctional proteins (27, 28, 62, 63) and we previously PMN transmigration triggered by HXA_3_ was required for barrier breach of pulmonary epithelium by *S. pneumoniae* (21, 31). PLY, which triggers epithelial production of HXA_3_ (20) and may enhance bacterial translocation. Indeed, a highly invasive *S. pneumoniae* serotype 1 strain, which releases high levels of PLY, induces severe inflammation of the lung, with ∼4-fold more neutrophils than a serotype 2 strain tested in parallel (14).

Although PLY clearly promotes PMN transmigration, monolayer infection with TIGR4*Δply* also triggered (lower levels of) neutrophil movement, presumably due to the production of chemoattractants in response to (non-PLY) bacterial products (64, 65). The level of barrier breach by TIGR4*Δply* in the presence of PMNs appeared elevated compared to background levels (Figures 1 and 2) but these higher levels did reach statistical significance. Interestingly, at a MOI of 20, PMN transmigration in response to *S. pneumoniae* infection resulted in the PLY-dependent loss of more than 50% of the epithelial monolayer, a loss that correlated with the efficient transepithelial movement of the marker HRP and the apparently unimpeded barrier breach by pneumococci. Epithelial detachment by activated neutrophils has been observed for several cell lines (66, 67) and has been correlated with secreted neutrophil elastase, which, during pneumococcal pneumonia, causes extensive tissue damage in both mouse models and humans (68, 69). Although dramatic epithelial loss provides an explanation for PLY- and PMN-promoted bacterial transmigration at a MOI of 20, at a lower MOI, wild type TIGR4 was capable of barrier breach in the absence of monolayer destruction, indicating that frank epithelial loss is not required for PLY-promoted bacterial movement in this experimental system.

In summary, by manipulating infectious dose and the presence or absence of PLY or PMNs during apical infection of polarize monolayers, we found that *S. pneumoniae* alters the epithelium and its interaction with PMNs in multiple ways. The bacterium utilizes PLY-dependent and PLY-independent mechanisms to induce cleavage and mislocalization of E-cadherin, as well to recruit PMNs across respiratory epithelial cells monolayers. Notably, E-cadherin disruption alone was insufficient to induce significant bacterial movement across epithelium, which instead depended on PMN transmigration. These studies provide further insight into the multifaceted role of PLY in pulmonary inflammation and barrier breach during *S. pneumoniae* lung infection.

## Acknowledgement

This work was supported by NIH Award Number 5R37AI037657-22 to RKT, NIGMS Award Number K12GM074869 Training in Education and Critical Research Skills (TEACRS) and 1SC2GM141988-01 to WA, and by the California State University Program for Education and Research in Biotechnology Graduate Student COVID-19 Research Restart Program (to DM).

## Supplemental Figure Legends

<u>Supplemental Figure 1: PLY-producing *S. pneumoniae* induces membrane permeabilization of epithelial cells and PMNs at high but not low dose infection.</u> **A.** Polarized monolayers of H292 cells on Transwells were infected with the indicated MOI of GFP-expressing WT or *Δply* SPN, and monolayer adhered SPN visualized by IF microscopy and bacteria adherence per area was quantitated. **B.** Polarized monolayers of H292 cells on Transwells were infected with the indicated MOI of WT SPN and stained with DAPI and PI, and percentage PI permeabilization was dertermined. **C.** H292 cells in suspension were infected with the indicated MOI of WT or *Δply* SPN, followed by flow cytometry analysis for PI incorporation. **D.** PMNs in suspension were infected with the indicated MOI of WT or *Δply* SPN, followed by flow cytometry analysis for PI incorporation. **E.** Released MPO activity in apical buffer of WT or *Δply* SPN infected monolayers post PMN transmigration and before lysis by detergent. Each panel shown is representative of three independent experiments in triplicates. Error bars represent mean ± SEM. Statistical analysis performed using One-way ANOVA: ** *P*-value ≤ 0.01, *** *P*-value ≤ 0.001, **** *P*-value ≤ 0.0001.

<u>Supplemental Figure 2: Single channel IF microscopy images of polarized H292 monolayers infected with the indicated MOI of WT *S. pneumoniae*.</u> Monolayers were stained with DAPI (blue), E-cadherin (green), and phalloidin (red). Scale bar = 20 μm.

<u>Supplemental Figure 3: PLY enhances MMP activity in culture supernatants of both epithelial cells and PMNs but MMP inhibition does not protect against junctional damage.</u> **A.** Polarized monolayers of H292 cells on Transwells were treated with vehicle (Veh.) or a wide-spectrum MMP inhibitor (GM6001) and intoxicated with 1U PLY as indicated, and MMP activity measured by fluorogenic peptide substrate conversion. **B.** PMNs in suspension were treated with vehicle or GM6001 and then intoxicated with 1U PLY as indicated, and MMP activity measured by fluorogenic peptide substrate conversion. **C-D.** Polarized monolayers of H292 cells on Transwells were treated with Veh. or GM6001 and intoxicated with 1U PLY or infected with WT SPN at MOI 2, as indicated. In **C.** monolayers were lysed for western blotting of E-cadherin. In **D.** monolayers were permeabilized and stained with E-cadherin antibody (green), DAPI (blue), and phalloidin (red). Scale bar = 20 μm. Peripheral E-cadherin organization was quantitated by image analysis with IJOQ. Each panel shown is representative of three independent experiments in triplicates. Error bars represent mean ± SEM. Statistical analysis performed using ordinary one-way ANOVA: *p-value ≤ 0.05, **p-value ≤ 0.01, ***p-value ≤ 0.001.

